# Divergent Thermal Reaction Norms of the Spontaneous Mutation Rate Among Populations of *Chironomus riparius*

**DOI:** 10.64898/2026.04.07.716879

**Authors:** Markus Pfenninger, Maria Esther Nieto-Blazquez, Burak Bulut

**Affiliations:** Department of Molecular Ecology, Senckenberg Biodiversity and Climate Research Centre, Georg-Voigt-Str. 14-16, D-60325, Frankfurt am Main, Germany; Institute for Molecular and Organismic Evolution, Johannes Gutenberg University, Johann-Joachim-Becker-Weg 7, D-55128, Mainz, Germany

**Keywords:** Population reaction norms, phenotypic plasticity, thermal regime, global change, Bayesian analysis

## Abstract

The germline mutation rate µ is a fundamental evolutionary parameter, yet its plasticity in response to environmental factors, particularly temperature, remains poorly understood. While often modeled as a species-specific constant, we tested whether µ evolves in response to local thermal regimes. Using whole-genome sequencing of mutation accumulation lines in the non-biting midge *Chironomus riparius*, we demonstrate divergent thermal reaction norms between populations from climatically distinct regions: Central Europe (Germany) and the Mediterranean (Spain). The Central European population displays a highly plastic, U-shaped reaction norm, whereas the Mediterranean population exhibits a more canalized, temperature-insensitive response. This divergence conforms to theoretical expectations: the higher thermal variance of high-latitude habitats selects for plasticity, while thermally more stable Mediterranean habitats favour robustness and optimises the mutational load in the respective thermal regimes. Furthermore, population-specific mutational spectra (Ts/Tv ratios) indicated evolved differences in DNA repair machinery. However, this is only partially mirrored by Reactive Oxygen Species (ROS) dynamics, where Mediterranean larvae maintain lower ROS levels and a buffered response to thermal extremes. These findings provide evidence for evolution of the mutation rate itself, challenging the assumption of constancy.

## Introduction

The germline mutation rate (µ) is one of the most important population genetic parameters, its value determining levels of genetic diversity, genetic load and responses to selection of populations and species (Beichman et al., 2024; Bertorelle et al., 2022; Carlson et al., 2020). Furthermore, accurate estimates of µ are essential for dating divergence in molecular phylogenetics (Tiley et al., 2020) or reconstruction of past demographic dynamics (Minin et al., 2008).

While µ has historically been regarded as a constant within species, several studies have shown that µ varies along the genome (Francioli et al., 2015; Gao et al., 2014; Nieto-Blazquez & Pfenninger, 2025), with age at reproduction (Bulut & Pfenninger, 2025; Pletcher et al., 1998; R. J. Wang et al., 2022) and exposure to anthropogenic substances (Bulut et al., 2024; Doria et al., 2021; Keith et al., 2021; Rigano et al., 2025). Most importantly for natural populations, µ exhibits plasticity in response to environmental parameters, particularly temperature (Bulut & Pfenninger, 2025; A.-M. Waldvogel & Pfenninger, 2021).

In the ectothermal non-biting midge *Chironomus riparius* the thermal reaction norm of µ is U-shaped. This curve results mainly from two opposing forces: generation time in ectothermic organisms like *C. riparius* is inherently linked to ambient temperature (Oppold et al., 2016). At low temperatures, the longer generation time will lead to a time-dependent mutation accumulation, increasing µ. Conversely, at high temperatures, the mutagenesis is governed by replication fidelity, which increases with shorter generation times (Bulut & Pfenninger, 2026). Consequently, µ can vary by a factor of four due to the seasonal changes of temperatures experienced by the midges throughout the year (A.-M. Waldvogel & Pfenninger, 2021). This implies that point estimates derived from standard laboratory temperatures may fail to reflect the effective mutation rates relevant to evolution in constantly fluctuating natural environments (Bell, 2010).

However, it remains unclear whether such thermal reaction norms are fixed within a species or if they vary between populations as a result of evolutionary processes. In general, evolutionary theory robustly predicts that phenotypic plasticity evolves to higher levels in environments that are variable but predictable, allowing organisms to match their phenotype to likely encountered selective pressures (Reed et al., 2010). In contrast, reduced plasticity should evolve in more constant environments, as plastic responses come with a cost (Lande, 2014). Specifically for mutation rates, the drift-barrier hypothesis or selection for genomic stability suggests that populations evolving in distinct thermal regimes should exhibit divergent reaction norms to minimize mutation load (Lynch, 2010). However, this has not yet been addressed in the literature. While differences in point estimates of µ among populations and closely related species have recently been shown for *Drosophila* (Y. Wang et al., 2023), the critical question remains whether the shape of the reaction norm evolves in response to the local environment. Understanding these reaction norms is vital e.g. for forecasting evolutionary responses to climate change (Oomen & Hutchings, 2022; A.-M. Waldvogel et al., 2020).

In this study, we explored the thermal reaction norms of spontaneous germline mutation rates along an environmentally realistic temperature gradient in *C. riparius* populations that evolved in distinct climatic zones in Central- (Hessen, Germany) and Mediterranean Europe (Andalusia, Spain) with accordingly different thermal regimes. The thermal regime in Andalusia allows for reproduction all year round at water temperatures ranging roughly from 12°C to 27°C (annual mean 19°C, (Lutz et al., 2016)). The Hessian population experiences signifcantly higher annual variation, ranging from below 4° during cold spells in winter to 22°C with a mean water temperature of 16°C during the main reproductive phase from April to October (A.-M. Waldvogel & Pfenninger, 2021). In addition to this annual pattern, substantial temporal and spatial small scale variation in temperature characterise the habitats of these midges (Leach et al., 2023). Previous studies have shown that the populations are overall adapted to their respective thermal regimes (Waldvogel et al., 2018). We further investigated *in vivo* Reactive Oxygen Species (ROS) levels across an overlapping temperature gradient to explore a potential proximate mechanism driving differences in thermal reaction norm.

## Material & Methods

### Populations assessed and rearing conditions

The *C. riparius* populations used in the present study are maintained as large (N > 1000) in-house laboratory cultures. One population originated in southern Spain (Arroyo del Pilar de la Dehesa, Andalusia; 37.399080°N, -4.5267980°E, sampled in 2013), the other from Germany (Hasselbach, Hessen, 50.167562°N, 9.083542°E, Figure 1A). While the Hessian population was started at the same time, but was regularly restocked from the original population, the Andalusian population remained isolated since initial sampling. Both stocks were meticulously kept apart to avoid their mixing.

**Figure 1.**
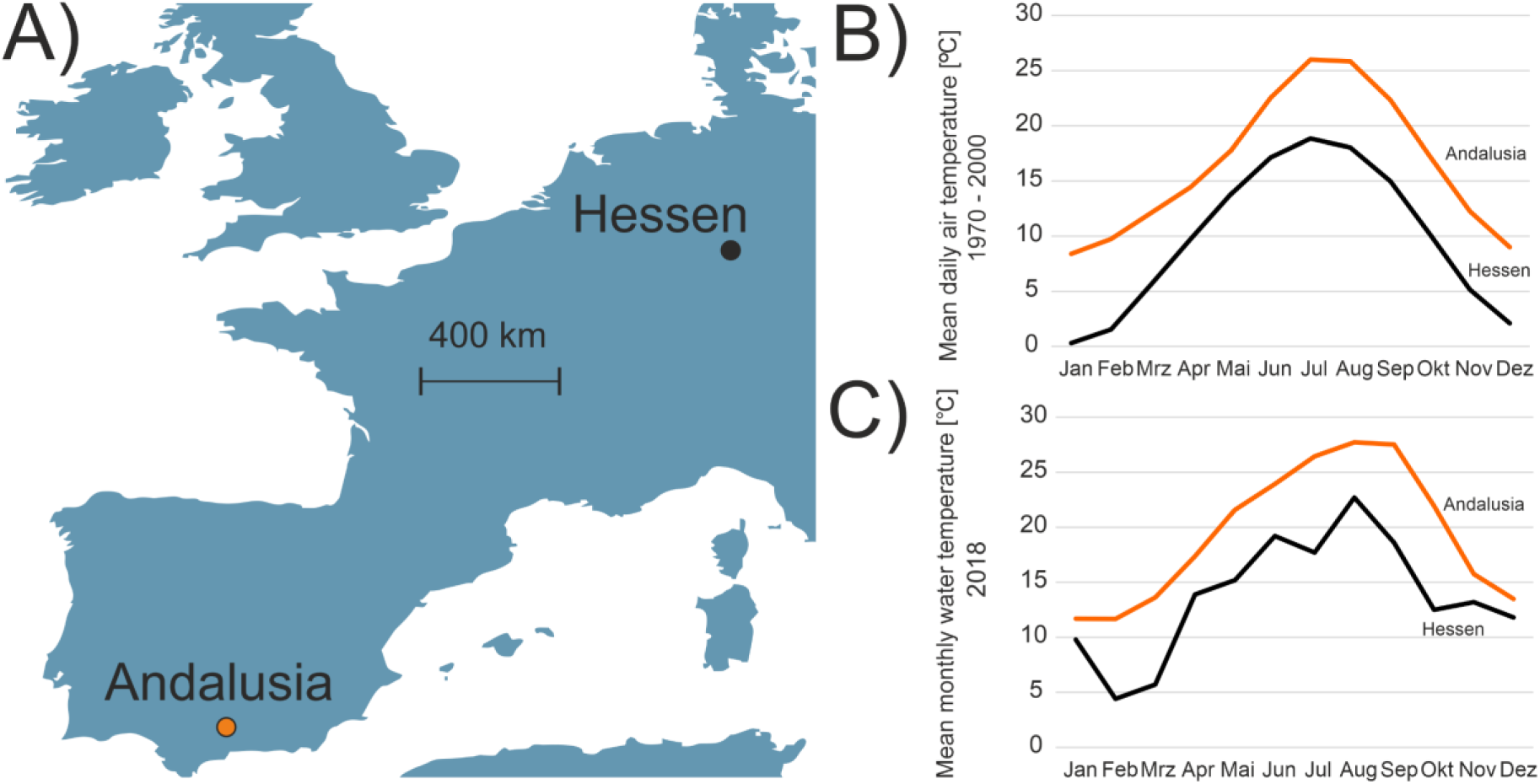
Geographical origin of populations and thermal regime at sampling sites. A) Origin of populations sampled in Europe. B) Mean daily air temperatures per month at the sampling sites according to Worldclim data for the period 1970 – 2000. C) Mean monthly stream temperatures in the area of origin in the year 2018.

Culturing conditions of both of the cultures follows a modified version of the method described in the OECD guideline N°219 and already published by (Foucault et al., 2019). Briefly, the cultures are continuously aerated and maintained at constant temperature of 20°C, 60% of humidity and under a 16:8 photoperiod. Larvae are raised in large trays with 1:4 ratio of sediment:medium. Sediment consists in washed (pH neutral) playground sand. Medium is deionised water adjusted to a conductivity of 520–540 μS/cm with aquarium sea salt (e.g., TropicMarin®) and a basic pH around 8. The organisms were fed daily with 0.4 g finely grounded fish food (e.g., Tetramin® Flakes).

The genome-wide F_ST_ among the two populations was estimated as 0.078 +/- 0.020 (A. Waldvogel et al., 2018). Both populations were obtained from natural but anthropogenically remodelled streams. Apart from the geographically and resulting different thermal regimes (Figure 1), these probably also differed in many other, selectively relevant aspects, such as water chemistry, water regime or emissions. All the research complies with applicable laws on sampling from natural populations and animal experimentation (including the ARRIVE guidelines).

### Mutation accumulation experiments

Because the species’ swarm-mating behaviour prevents direct identification of the (usually monandrous) parents, their allelic composition is instead reconstructed from pooled sequencing of their immediate offspring (Oppold & Pfenninger, 2017). To achieve this, a single egg-mass is brought to hatch under controlled conditions, the full siblings develop to adults, mate and reproduce (Foucault et al. 2019), thus starting a Mutation Accumulation Line (MAL). A single egg-mass from the peak reproduction period is then randomly chosen for the next generation. The remaining siblings (several hundred) from this first generation of each MAL are pooled to generate a “parental pool”. While this pooling approach with a large number of offspring does not allow to reconstruct the parents’ genotypes, it is a robust method for accurately reconstructing the parental allele composition at each locus, with potential biases mitigated by high sequencing depth (50X) and stringent variant filtering criteria (Oppold & Pfenninger, 2017, see Supplemental Figure). We established MALs at five temperatures: 12°C, 14°C, 17°C, 23°C, and 26°C. Ten MALs were initiated for each temperature condition, with additional backup MALs to compensate for the loss of lines due to lethal inbreeding effects.

### Whole genome sequencing and bioinformatics

DNA was extracted using the DNeasy Blood and Tissue Kit (QIAGEN). To establish an ancestral baseline for identifying *de novo* mutations (DNMs), we sequenced a pool consisting of one leg from each of 120 F1 individuals (60× target coverage). After five generations, one female from each MA-line was sequenced on the Illumina NovaSeq 6000 platform (30× target coverage) following standard library preparation. Data processing followed GATK best practices (Pettrich et al., 2025)McKenna et al., 2010). Reads were paired using PEAR (Zhang et al., 2014) and mapped to the *C. riparius* reference genome v.4 (Pettrich et al., 2025) using BWA-MEM. Duplicates were marked with Picard v.1.123, and low-quality reads were removed using SAMtools (Li et al., 2009). Local realignment and base recalibration were performed using GATK. BAM files were merged, and DNMs were identified using accuMUlate (Winter et al., 2018). Candidate mutations were filtered using a custom script with the following thresholds: mutation probability > 0.90; correct descendant genotype probability > 0.90; zero mutant reads in the ancestor; mapping quality difference < 2.95; and strand bias > 0.05. All filtered sites were manually validated using IGV. The mutation rate µ was calculated as the total number of confirmed mutations divided by the product of callable sites and the number of generations.

### Statistical analysis

The best fit model for the thermal reaction norms was determined using AIC (Akaike, 2003). To investigate differences in phenotypic population reaction norms (PRNs) of µ, respectively ROS to temperature, we employed a Bayesian hierarchical modelling framework using the *brms* package (v. 2.22.0) in R (v. 4.2.2), which interfaces with Stan for Hamiltonian Monte Carlo (HMC) sampling (Bürkner, 2021). Parameters were estimated using the No-U-Turn Sampler (NUTS). We ran 4 independent Markov chains for 20,000 iterations each, with a warm-up period of 3,000 iterations, resulting in 68,000 total post-warmup posterior draws. For the fixed effect coefficients, we used flat prior distributions. The population intercept and residual variance were assigned Student_t priors (3, 6.5, 3.7) and (3, 0, 3.7), respectively, following *brms* defaults. Model convergence was assessed using the ^R diagnostic and Effective Sample Size (ESS). All parameters reached convergence with ^R < 1.01 and both Bulk and Tail ESS > 45,000. Unlike frequentist approaches that rely on non-overlapping confidence intervals at specific temperatures, our Bayesian model assesses the probability that the functional form (shape) of the reaction norm differs. High posterior probabilities of non-zero interaction terms provide robust evidence for divergent evolution, even in the presence of (expected) individual variation.

Mutation rates of the two populations at 15°C, respectively 26°C were compared with a Bayesian implementation of a Poisson-test. The observed overall Ts/Tv rates of the two populations were compared with a Bayesian test of proportions. We obtained genome-wide Watterson’s theta (ϴ) estimates with PoPoolation1 v.1.2.2 (Kofler et al., 2011) using 10kb windows for field-obtained PoolSeq samples (A. Waldvogel et al., 2018) of the two populations were compared with a Bayesian t-test. All tests were performed with the R package *BayesianFirstAid* v. 0.1 (Bååth, 2014), using default values.

### ROS Measurement

L3-stage larvae were used in this analysis because of their physiological robustness. We quantified ROS using CellROX Orange (Thermo Fisher), a cell-permeable probe that exhibits fluorescence at 545/565 nm upon oxidation by various ROS (e.g., hydrogen peroxide, hydroxyl radical). This method allows for accurate relative estimation of overall ROS levels in living organisms (Kang et al., 2013; Wu et al., 2019).

Single larvae (n = 20 per treatment) were placed in 24-well plates with 2.5 mL medium, allowing sufficient passive oxygen diffusion to prevent hypoxia. Larvae were stained with 1.5 µM CellROX Orange, a concentration optimized to provide detectable signals without toxicity. Larvae were exposed for 24 hours to seven temperature regimes (4°C to 28°C in 4°C increments) in a climate chamber (550 lux; 16:8 light/dark cycle). This duration was chosen to capture the acute oxidative stress response. Random sampling from an asynchronous population minimized systematic variation.

Following treatment, plates were insulated during in-house transport to prevent temperature shocks. Live imaging was performed using a ZEISS Axio Imager 2 (10x magnification) with a 1s exposure time and a "43 HE" filter set (excitation ∼550 nm; emission >570 nm). Fluorescence was consistently quantified in the first abdominal segment (Figure A2).

Images were analyzed using ImageJ Fiji (v. 2.15.0). Images were converted to 8-bit grayscale and thresholds were adjusted to minimize background noise resulting from high-intensity illumination. Mean fluorescence intensity served as a measure of relative ROS production. While not an absolute quantification, the uniform application of optimized reagent concentrations across all treatments ensured valid relative comparisons of in vivo ROS dynamics.

## Results

### Mutation rate reaction norms and -spectra are population specific

A list with all observed mutations, their positions in the genome and the calling statistics can be found in Supplemental Table 1. Due to issues with keeping temperature stable in the 23°C experiment with the Hessian population, results from this temperature are entirely missing.

The thermal reaction norm of the Hessian population was best described by a 2nd order polynomial function (ΔAIC > 50 compared to a linear model; Figure 2A), confirming the U-shaped response previously described (Waldvogel & Pfenninger, 2021). In contrast, the Andalusian population showed a distinct lack of thermal sensitivity; the polynomial model offered only a marginal improvement over a linear model ΔAIC = 7), and a linear model with zero slope provided a good fit, suggesting a canalized mutation rate across temperatures.

**Figure 2.**
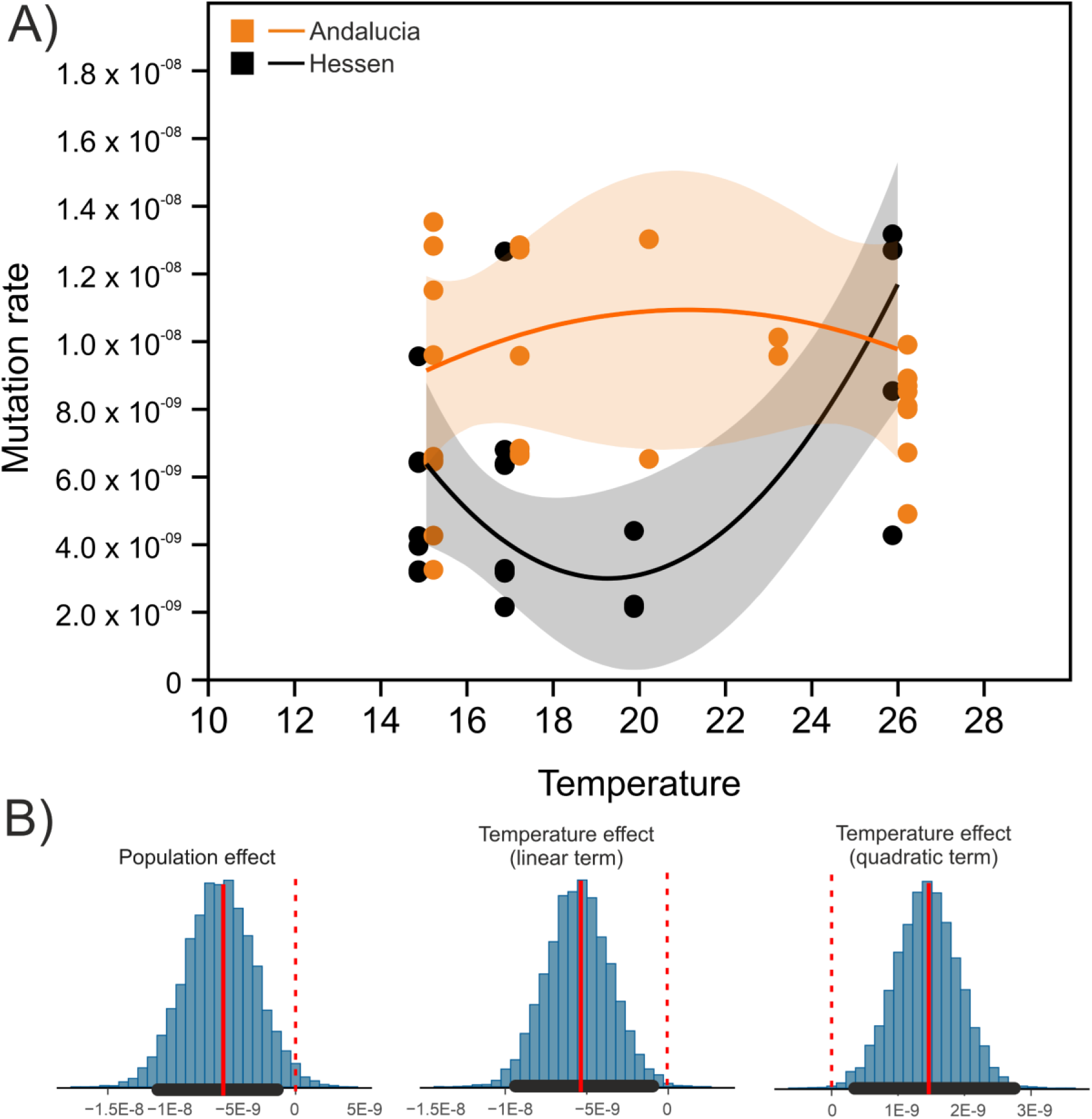
Effect of natural temperature gradient on germline mutation rate. A) Per stMAL mutation rates of two populations (orange: Andalusia, black: Hessen) to temperature. The fitted lines correspond to a 2^nd^ order polynomial function, the shades correspond to the . B) Posterior distributions of effects and parameters on µ in Bayesian analysis. Y-axes are frequencies of parameter values sampled from the posterior distribution. Red vertical line indicates the median estimate, red dashed lines are the zero effect line, the horizontal bold black bar the 95% HDI. All effects had a posterior probability of > 95% to be different from zero. Y-axes are frequencies of sampled parameters in Monte-Carlo Markov Chains runs of posterior distributions.

Bayesian analysis supported this divergence in reaction norm shape. The interaction terms for both quadratic and linear parameters were distinct from zero with high certainty (pp = 1.0 and 0.99, respectively), confirming that the shape of the reaction norms differs significantly between populations (Figure 2B). Examining the plot of the reaction norms with the associated 95% confidence intervals showed a significant difference in the range of ∼15.5 – 22°C. Overall, the Hessian population exhibited a lower mutation rate across the temperature range (median reduction of µ = -6.02 x 10^-8^ (95% HDI -1.15 x 10^-7^ - -4.66 x 10^-9^, Figure 2B).

When examining specific thermal points, there was moderately strong support for the Hessian population having a 30% lower µ at 15°C (pp = 88.3%, 95% HDI 0.41 – 1.3; Figure 3A). This pattern inverted at high temperatures (26°C), where the Andalusian population showed a 17% lower mutation rate (pp = 75.4%, 95% HDI 0.69 – 2.2; Figure 3B). The mutational spectrum showed strong differentiation: the Transition/Transversion (Ts/Tv) ratio was 0.59 (95% HDI 0.43 – 0.75) for Hessen and 0.41 (95% HDI 0.29 – 0.54) for Andalusia (pp difference = 94.9%; Figure 3C).

**Figure 3.**
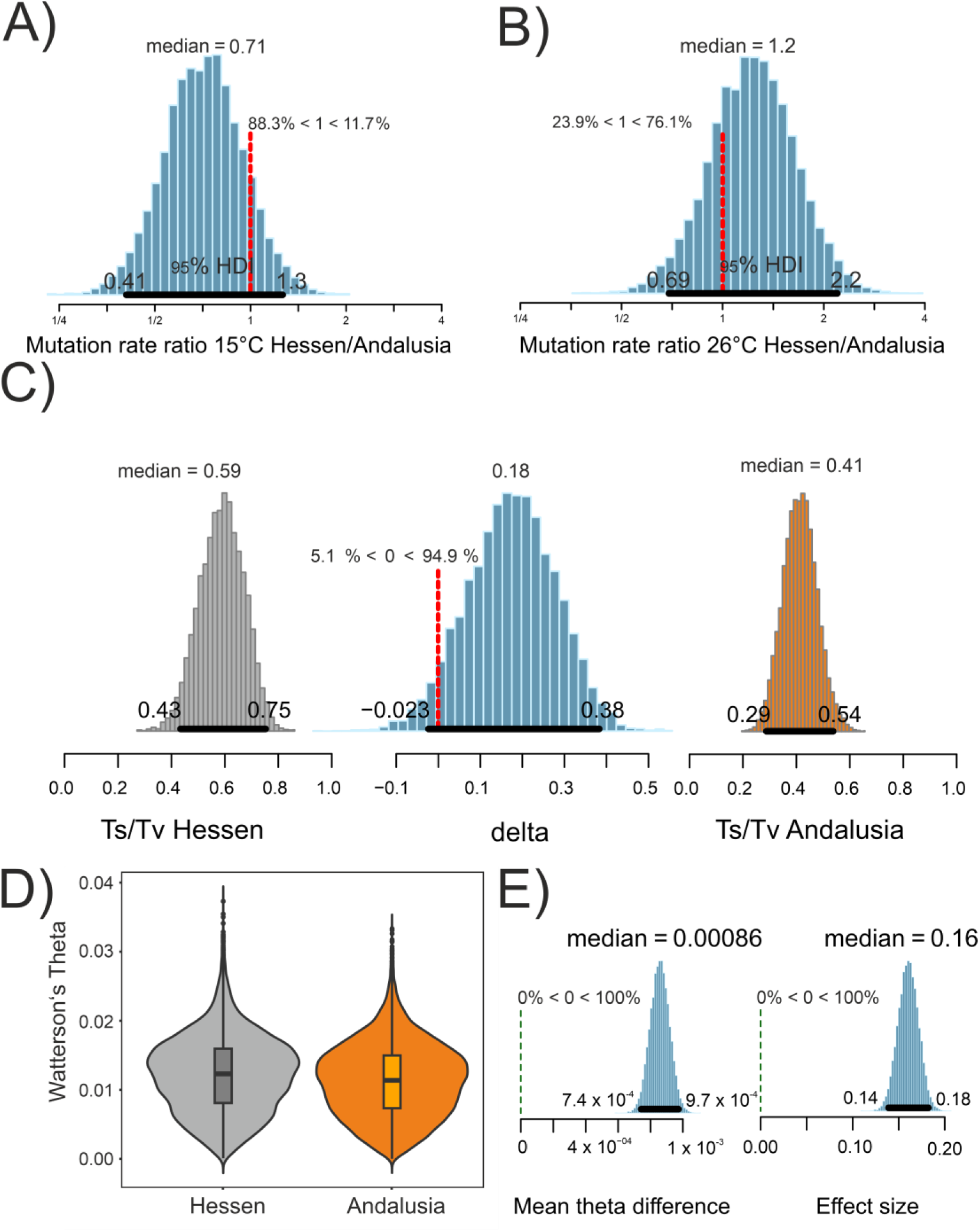
Evidence for differential evolution. A) Posterior distribution of mutation rate ratios Hessen/Andalusia at 15°C. B) The same for 26°C. C) Bayesian comparison of the Transition/Transversion ratio. D) Violin plot of the genome-wide estimates of Watterson’s theta for Hessen and Andalusia . E) Bayesian paired t-test of theta per genome window. If not otherwise stated, Y-axes are frequencies of sampled parameters in Monte-Carlo Markov Chains runs of posterior distributions.

Genome-wide ϴ estimates were significantly higher in the natural population in Hessen (0.0122; 95% HDI 0.0121 – 0.0123) compared to Andalusia (0.0113; 95% HDI 0.0112 – 0.0114; Figure 3D). While the probability of this difference was maximal (pp = 100%), the effect size was negligible (Cohen’s D = 0.160; Figure 3E).

We used the inferred thermal reaction norms of µ to calculate the expected mutation rates for the mean monthly water temperatures in the respective habitats (Figure 4A). We then computed the expected number of generations per month (Figure 4A), based on the known relation between water temperature and generation time (Oppold et al., 2016). This allowed to estimate an effective mean µ over the respective reproductive seasons (5.71 x 10^-9^ in Hessen and 8.02 x 10^-9^ in Andalusia, Figure 4B). The difference in effective µ is thus 2.31 x 10^-9^ among the populations. Applying the reaction norm of the respectively other population to the local thermal regimes showed that the expected effective mutation rate would in increase in both populations (plus 3.8 x 10^-9^ in Hessen, plus 2.0 x 10^-10^ in Andalusia, Figure 4B).

**Figure 4.**
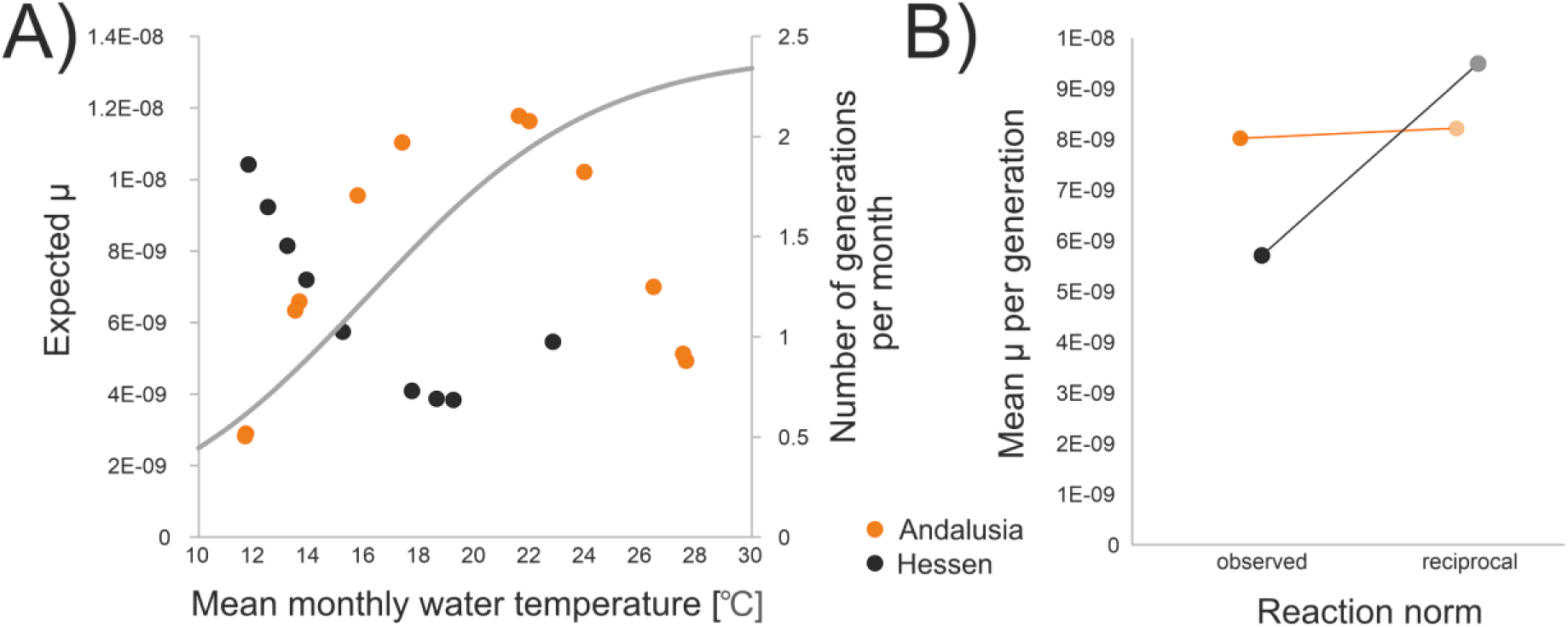
Effective mean mutation rates. A) Expected µ according to thermal reaction norms for the mean monthly water temperatures during the reproductive phase in Andalusia (orange dots) and Hessen (black dots). The grey curve indicates the expected number of generations per month for the temperature range. B) Estimated effective mean µ per generation for the observed reaction norm (left half) and for the reaction norm of the respectively other population (right half, light coloured dots), given the observed thermal regime.

### Population specific thermal reaction norms of ROS production

The impact of temperature on *in vivo* ROS concentration was best described by a 2nd order polynomial function in both populations (ΔAIC > 200 compared to linear models; Figure 5A). The magnitude of ROS production differed substantially. Bayesian analysis indicated with maximal support (pp = 100%) that the Andalusian population maintained lower ROS levels overall (median fluorescence reduction = - 52.95; 95% HDI -33.92 to -73.17; Figure 5B). Furthermore, the reaction norm shape differed; the Andalusian population exhibited a significantly less steep increase in ROS at extreme temperatures, evidenced by interaction terms for both quadratic and linear parameters differing from zero with highest certainty (Figure 5B). The physiological thermal optima (nadirs of the fitted functions) were statistically indistinguishable between populations (Hessen: 14.87°C, 95% CI 10.85–20.22; Andalusia: 13.48°C, 95% CI 7.90–22.56).

**Figure 5.**
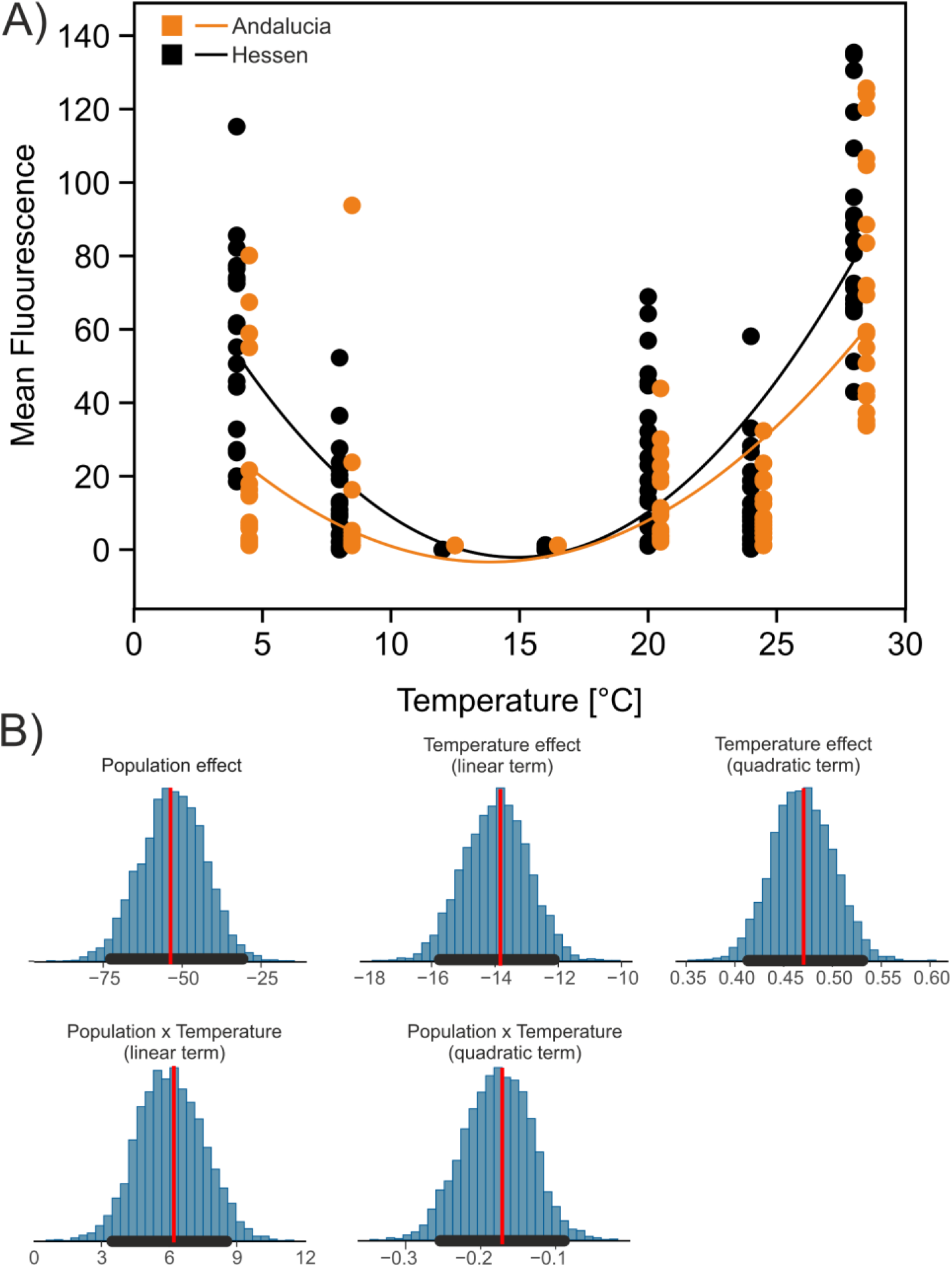
Reaction norm of *in vivo* ROS levels along a natural temperature gradient. A) Mean fluorescence proportional to the presence of ROS of individuals of two populations (orange: Andalusia, black: Hessen) to temperature. The fitted lines correspond to a 2^nd^ order polynomial function. B) Posterior distributions of effects and parameters on the µ in Bayesian analysis. Red vertical line indicates the median estimate, the horizontal bold black bar the 95% HDI. All effects are with highest certainty different from zero (pp = 100%). Y-axes are frequencies of sampled parameters in Monte-Carlo Markov Chains runs of posterior distributions.

## Discussion

### Divergent Evolution of Mutation Rate Reaction Norms

Our results demonstrate that the thermal reaction norm of µ is not a fixed species-specific trait but has evolved divergently between populations. While the Hessian population exhibits the U-shaped reaction norm previously described for *Chironomus riparius*, characterized by high plasticity (Waldvogel & Pfenninger, 2021), the Andalusian population displays a significantly flatter, canalised response.

In principle, the differential reaction norms could have arisen due to the different rearing histories of the laboratory populations used in the experiment starting in 2023. Both populations were established at the same time in the laboratory (2013), but contrary to the Hessian population that was regularly refreshed from the original population, the Andalusian population was kept isolated in the laboratory at constant benign conditions since the initial sampling. Even though the population was kept at high densities, and it has thus experienced a substantial amount of genetic drift. However, obtaining the observed pattern by drift would imply that specifically alleles responsible for a plastic reaction would have been lost in the Andalusian lab population. Since plasticity is most likely itself a polygenic trait (Kovuri et al., 2023), such a directed random loss at several loci simultaneously appears rather unlikely. The generally benign constant conditions in the laboratory have certainly relaxed the natural selection on many traits (Hoffmann & Ross, 2018), a regime to which the Andalusian population was longer exposed than the regularly refreshed Hessian population. There is no clear expectation whether relaxed selection should actually lead to a loss of plasticity. On the one hand, plasticity of traits that are not subject to direct selection may actually allow plasticity to persist longer (Lahti et al., 2009). On the other hand, if the costs of maintaining plasticity are high, selection is expected to reduce plasticity in constant environments (Gomez-Mestre & Jovani, 2013).

However, it is not clear in the current case whether plasticity needed to evolve and therefore requires costly stabilising selection to maintain it – or whether it is rather canalisation that evolved (Sommer, 2020). By experimentally separating the effect of temperature and generation time, the study of (Bulut & Pfenninger, 2026) suggested that the observed U-shaped thermal plasticity of µ in *C. riparius* is mainly the mechanistic result of two opposing forces: decreasing replication fidelity when the generation time becomes shorter and time dependent accumulation of mutations with increasing generation time, with an optimum in between. As generation time in natural populations of the ectothermic midges is strongly dependent on ambient temperature, the U-shaped plastic thermal reaction norm would thus be a logical, natural outcome of these two processes, requiring rather the observed canalisation in the Andalusian population to evolve and be maintained.

Laboratory conditions are not unequivocally beneficial and may trigger adaptation to this specific environment (Pfenninger & Foucault, 2020). Selection of µ to the more or less constant room temperature therefore cannot be excluded for the Andalusian population which was longer exposed to it. An adaptation to the actually experienced constant conditions does not explain the evolution of the reaction norm to a temperature gradient, which would either require to experience thermal variability or that optimisation to a particular temperature is unavoidably linked to the loss of plasticity (Scheiner & Levis, 2021). However, an adaptation to room temperature appears unlikely, because µ at 20°C is not the lowest value for the Andalusian population.

While the above considerations render a laboratory origin of the observed differences less likely, there are several natural processes that could have led to different reaction norms in the two populations. Given the high effective population size of natural *C. riparius* populations (A. Waldvogel et al., 2018), leading to the propensity of rapid adaptation even at moderate selective pressures (Rigano et al., 2026) and the substantial fitness relevance of the trait (Lynch et al., 2016), it appears highly unlikely that the thermal reaction norms of the natural populations diverged by random genetic drift. Also, a difference in selective efficiency among the populations is unlikely as theta, the mutation-population parameter is only marginally different among the populations (delta theta = 0.00085).

Differences in the thermal regime exerting selective pressure in either one (conditional neutrality) or divergently in both populations (local adaptation) could have driven the observed difference in reaction norm. The annual temperature range experienced in Germany is about three degrees Celsius larger than in Andalusia (18°C vs. 15°C) and closer to the minimal temperature required for reproduction. While this difference may appear negligible e.g. in a vertebrate context, it is substantial for ectothermic organisms (Paaijmans et al., 2013; Scranton & Amarasekare, 2017) and even a 3°C range difference can significantly impact physiological performance e.g. during reproduction (Massey & Hutchings, 2021). The different thermal regimes therefore likely exert substantially different selection pressures on the two populations.

There is evidence that the different reaction norms minimise mutational load under the respective thermal regimes. We observed a "crossing of lines" at thermal extremes: the Hessian population exhibits a lower µ at 15°C, a temperature frequently encountered in its habitat, while the Andalusian population shows a lower µ at 26°C, with, admittedly, limited statistical support. More importantly, calculating an effective average µ for the respective thermal regimes in the two populations showed that the observed reaction norms minimise the mutational load relative to the respectively other reaction norm. In other words, tentatively equating a lower mutational load with higher fitness, the respective reaction norms increase the fitness of the populations given the local thermal regimes, albeit this effect was much larger for the Hessian than the Andalusian population. While definitive proof for reciprocal local adaptation of the thermal reaction norm of µ would need to be experimentally established, it appears that there is at least a good fit between the respective environmental variability and the plasticity of the trait. Interestingly, the total mutational input is quite similar for both populations (difference of average effective µ = 2.3 x 10^-9^), despite the much larger mean difference among reaction norms (6.02 x 10^-8^), likely leading to the observed similar theta distributions (assuming historically large and constant N_e_). The observed divergence aligns with theoretical expectations regarding environmental stability, more plasticity in the more variable environment and vice versa. This supports the view that mutation rates are not merely passive physiological consequences but are actively shaped by selection to minimise genetic load under local environmental conditions. Notably, this differentiation in reaction norms persists despite the species generally exhibiting low population differentiation due to high gene flow in a stepping stone structure (A. Waldvogel et al., 2018), highlighting the strength of local selection pressures.

### Proximate Mechanisms: Oxidative Stress and DNA Repair

The physiological data suggest that differential regulation of Reactive Oxygen Species (ROS) has limited influence on the observed divergence in mutation rates. Besides studies showing the association of ROS with increased mutation rates in an ecotoxicological context (Bulut et al., 2024; Rigano et al., 2025), it has been mechanistically proven that ROS (in particular 8-oxoguanine) causes DNA damage leading to germline mutations (Cordero et al., 2024; Ohno et al., 2014).

The Andalusian population showed lower ROS levels over the entire temperature range tested and a less steep increase in ROS production at extreme temperatures. This implies on the one hand that the southern population has evolved more efficient oxidative stress protection mechanisms, likely an adaptation to the generally higher metabolic rates and thermal stress associated with their warmer habitat. On the other hand, however, the overall higher mutation rate in this population suggested that the germline mutation rate is driven by other processes. This aligns with other studies that show limited effects of thermal oxidative stress on the germline mutation rate (Berger et al., 2017; Joyner-Matos et al., 2011; Rajaei et al., 2021).

The significant difference in the transition/transversion (Ts/Tv) ratio between populations points to divergence in the molecular machinery itself. Since DNA damages that ultimately lead to transitions or transversions are often repaired by different pathways (Cebrián-Sastre et al., 2021; Pilzecker & Jacobs, 2019), this shift in mutational spectrum suggests that the populations differ not just in ROS prevention, but potentially in the efficacy or usage of specific DNA repair pathways. The interplay of demography and mechanism is also evident: the Hessian population, facing longer phases of lower mean temperatures and therefore longer, more variable generation times (Oppold et al., 2016), likely accumulates mutations via time-dependent mechanisms, whereas the Andalusian population faces primarily metabolism-dependent (thermal stress) mutagenesis.

### Implications for Population Genetics and Phylogenetics

The demonstrated spatial and environmental volatility of µ has profound consequences for evolutionary inference. First, the assumed mutation rates measured under benign laboratory conditions may be too low to reflect the realised mutation rate in nature. Consequently, divergence time estimates in phylogenetics that rely on a static clock rate may be substantially biased (Soni et al., 2024). Second, standard methods for molecular dating and coalescence analysis typically assume that µ is constant over time and space. Our results show that this assumption is invalid for species in fluctuating environments. Specifically, coalescence approaches (e.g., PSMC, MSMC) estimate the product of N_e_ x µ (Mather et al., 2020). If µ differs significantly and heritably between populations, this may be as well true for environmental changes over time (e.g. the Pleistocene climatic variability). Inferred changes in effective population size N_e_ may therefore actually reflect plastic or evolutionary shifts in µ or the intricated dependency of µ on changes in N_e_ (Lynch et al., 2016).

### Conclusion and Future Directions

We demonstrated that the spontaneous mutation rate is not a static species-specific parameter but rather a normal trait that possesses a thermal reaction norm potentially evolving in response to local climatic conditions. These reaction norms were significantly different for two evolutionarily relevant traits (mutation rate and ROS production), contributing to the active debate on climate-related reaction norm evolution (Oomen & Hutchings, 2022). While some studies find evidence for such evolution (Delgado et al., 2020; Kruger et al., 2022; Terblanche et al., 2009), others do not (Clemson et al., 2016; Matesanz et al., 2020). To further understand the dynamics of this evolution, future research should address three key areas. First, the scope of divergence. Do other fitness-related traits (e.g., metabolic rate, developmental timing, gene expression) exhibit similar divergence in thermal reaction norms? Second, questions on the evolutionary predictability. Along a climate gradient, is the change in reaction norm continuous and predictable, or does it exhibit tipping points and idiosyncratic behaviour? And third, the genetic basis of reaction norms. Can the population-specific reaction norms be predicted from underlying gene expression patterns, and which regulatory sequences drive these differences?

## Acknowledgments

We gratefully acknowledge the support of this study by a grant of the Deutsche Forschungsgemeinschaft (PF 390/15-1) to MP.

## Data Availability

All nucleotide sequences used in this study are deposited in the European Nucleotide Archive (ENA) under the project number PRJEB101187. Raw data for the analyses were made available at Zenodo doi: 10.5281/zenodo.17879081.

## Declaration of generative AI and AI-assisted technologies in the writing process

During the preparation of this work the authors used Gemini (Google, accessed October - December 2025) in order to improve the readability and language of the manuscript. After using this service, the authors reviewed and edited the content as needed and take full responsibility for the content of the published article.

## CRediT author statement

Markus Pfenninger: Conceptualization, Methodology, Validation, Resources, Project administration, Writing – original draft, Funding acquisition; Maria Esther Nieto-Blazquez: Writing – review and editing; Burak Bulut: Data curation, Formal analysis, Methodology, Writing – review and editing.

