## Supplementary Info for "Divergent Thermal Reaction Norms of the Spontaneous Mutation Rate Among Populations of *Chironomus riparius*" (from another version of this preprint: version 1, 2026-04-09)

1. Supplemental Table 1: List of mutations observed
2. Supplemental Figure 1: Estimated reaction norms with associated 95% confidence intervals

Supplemental Table 1. List of mutations.

| Population | °C | Chromosome | Bed interval start | Bed interval end | Ref allele | Mutant sample | Mutation direction | Probability of mutation | Probability of one mutation | Probability of correct descentant genotype | Likelihood |
|---|---|---|---|---|---|---|---|---|---|---|---|
| **Hessen** | 15 | Chr3 | 6030562 | 6030563 | G | H15 2 | GG->AG | 0.999209 | 0.999209 | 1 | 1.3795e-10 |
| **Hessen** | 15 | Chr3 | 14270075 | 14270076 | C | H15 22 | CC->AC | 0.999642 | 0.999642 | 1 | 1.3789e-10 |
| **Hessen** | 15 | Chr3 | 35762282 | 35762283 | C | H15 22 | CC->CG | 1 | 1 | 1 | 1.37841e-10 |
| **Hessen** | 15 | Chr2 | 31959569 | 31959570 | C | H15 23 | CC->AA | 1 | 1 | 0.999999 | 1.73893e-17 |
| **Hessen** | 15 | Chr2 | 50634499 | 50634500 | T | H15 23 | TT->AA | 0.903972 | 0.903972 | 0.902162 | 1.41808e-11 |
| **Hessen** | 15 | Chr2 | 51879527 | 51879528 | T | H15 23 | TT->CC | 0.999799 | 0.999799 | 0.983829 | 1.94325e-12 |
| **Hessen** | 15 | Chr2 | 54677789 | 54677790 | T | H15 23 | AA->TT | 0.999863 | 0.999863 | 0.991067 | 1.82646e-16 |
| **Hessen** | 15 | Chr2 | 55227197 | 55227198 | A | H15 23 | AA->GG | 0.903972 | 0.903972 | 0.902162 | 1.41808e-11 |
| **Hessen** | 15 | Chr2 | 55291346 | 55291347 | C | H15 23 | CC->AA | 0.974464 | 0.974464 | 0.945767 | 1.35189e-12 |
| **Hessen** | 15 | Chr2 | 55291349 | 55291350 | T | H15 23 | TT->AA | 0.988578 | 0.988578 | 0.945767 | 6.85645e-12 |
| **Hessen** | 15 | Chr2 | 55291352 | 55291353 | A | H15 23 | AA->GG | 0.988578 | 0.988578 | 0.945767 | 6.85645e-12 |
| **Hessen** | 15 | Chr3 | 21050490 | 21050491 | G | H15 23 | GG->AA | 0.974464 | 0.974464 | 0.945767 | 1.35189e-12 |
| **Hessen** | 15 | Chr2 | 8760375 | 8760376 | G | H15 26 | GG->AG | 0.999817 | 0.999817 | 1 | 1.37866e-10 |
| **Hessen** | 15 | Chr2 | 13230796 | 13230797 | C | H15 26 | CC->CT | 1 | 1 | 0.999999 | 1.37841e-10 |
| **Hessen** | 15 | Chr1 | 23840114 | 23840115 | C | H15 27 | CC->CT | 0.999976 | 0.999976 | 1 | 1.37844e-10 |
| **Hessen** | 15 | Chr1 | 49638138 | 49638139 | T | H15 28 | TT->CT | 1 | 1 | 1 | 7.0922e-10 |
| **Hessen** | 15 | Chr2 | 9803189 | 9803190 | C | H15 28 | CC->CT | 0.999217 | 0.999217 | 1 | 1.37949e-10 |
| **Hessen** | 15 | Chr2 | 30425988 | 30425989 | A | H15 28 | AA->AT | 0.977421 | 0.977421 | 0.965032 | 7.25604e-10 |
| **Hessen** | 15 | Chr1 | 48601281 | 48601282 | G | H15 29 | GG->AG | 0.999968 | 0.999968 | 1 | 1.37846e-10 |
| **Hessen** | 15 | Scaffold1 | 355256 | 355257 | A | H15 29 | AA->GG | 0.903972 | 0.903972 | 0.902162 | 1.41808e-11 |
| **Hessen** | 15 | Chr1 | 47632363 | 47632364 | T | H15 3 | TT->CT | 0.99882 | 0.99882 | 1 | 7.10058e-10 |
| **Hessen** | 15 | Chr3 | 42964695 | 42964696 | G | H15 3 | GG->CG | 0.995158 | 0.995158 | 1 | 1.38512e-10 |
| **Hessen** | 15 | Chr3 | 12693953 | 12693954 | A | H15 5 | AA->AT | 0.99794 | 0.99794 | 1 | 7.10685e-10 |
| **Hessen** | 17 | Chr2 | 25166303 | 25166304 | G | H17 1 | GG->AG | 0.999997 | 0.999997 | 1 | 1.37841e-10 |
| **Hessen** | 17 | Chr1 | 3276837 | 3276838 | C | H17 10 | CC->CT | 0.998636 | 0.998636 | 1 | 1.38029e-10 |
| **Hessen** | 17 | Chr2 | 2628596 | 2628597 | T | H17 12 | TT->AT | 0.991223 | 0.991223 | 1 | 7.155e-10 |
| **Hessen** | 17 | Chr4 | 92015 | 92016 | C | H17 12 | CC->CT | 0.998636 | 0.998636 | 1 | 1.38029e-10 |
| **Hessen** | 17 | Chr2 | 23601553 | 23601554 | T | H17 21 | TT->AT | 0.998821 | 0.998821 | 1 | 7.10057e-10 |
| **Hessen** | 17 | Chr4 | 6876221 | 6876222 | A | H17 21 | AA->AG | 0.996759 | 0.996759 | 1 | 7.11527e-10 |
| **Hessen** | 17 | Chr3 | 5834973 | 5834974 | A | H17 23 | AA->AC | 0.963161 | 0.963161 | 1 | 7.36347e-10 |
| **Hessen** | 17 | Chr1 | 14383814 | 14383815 | A | H17 25 | AA->AG | 0.999996 | 0.999991 | 1 | 1.26232e-13 |
| **Hessen** | 17 | Chr3 | 455166 | 455167 | C | H17 25 | CC->CT | 0.999217 | 0.999217 | 1 | 1.37949e-10 |
| **Hessen** | 17 | Chr2 | 45062381 | 45062382 | A | H17 28 | TT->AT | 0.977408 | 0.977408 | 0.965019 | 1.25872e-13 |
| **Hessen** | 17 | Chr3 | 26724219 | 26724220 | T | H17 28 | TT->AT | 0.977877 | 0.977877 | 1 | 7.25265e-10 |
| **Hessen** | 17 | Chr4 | 4088444 | 4088445 | G | H17 29 | GG->AG | 0.999976 | 0.999976 | 1 | 1.37844e-10 |
| **Hessen** | 17 | Chr3 | 46772649 | 46772650 | G | H17 30 | GG->AG | 0.990417 | 0.990417 | 1 | 1.39175e-10 |
| **Hessen** | 17 | Chr3 | 707708 | 707709 | C | H17 7 | CC->CG | 1 | 1 | 1 | 1.37841e-10 |
| **Hessen** | 17 | Chr1 | 15401030 | 15401031 | G | H17 9 | GG->AG | 0.996469 | 0.996469 | 1 | 1.38329e-10 |
| **Hessen** | 17 | Chr1 | 26400766 | 26400767 | A | H17 9 | AA->GG | 1 | 1 | 0.998519 | 1.75305e-13 |

| | | | | | | | | | | | |
|---|---|---|---|---|---|---|---|---|---|---|---|
| **Hessen** | 17 | Chr1 | 51248423 | 51248424 | C | H17 9 | CC->CT | 0.999048 | 0.999048 | 1 | 1.37972e-10 |
| **Hessen** | 17 | Chr2 | 6760391 | 6760392 | T | H17 9 | TT->GT | 0.999998 | 0.999998 | 1 | 7.09222e-10 |
| **Hessen** | 20 | Chr1 | 19733843 | 19733844 | A | H20 10 | AA->AG | 0.999999 | 0.999999 | 1 | 7.09221e-10 |
| **Hessen** | 20 | Chr4 | 7274951 | 7274952 | A | H20 12 | AA->AG | 0.999651 | 0.999651 | 1 | 7.09468e-10 |
| **Hessen** | 20 | Chr2 | 1982817 | 1982818 | G | H20 4 | GG->CG | 0.995678 | 0.995678 | 1 | 1.38439e-10 |
| **Hessen** | 20 | Chr2 | 23848692 | 23848693 | C | H20 8 | CC->CT | 0.999476 | 0.999476 | 1 | 1.37913e-10 |
| **Hessen** | 20 | Chr4 | 7484689 | 7484690 | G | H20 8 | GG->GT | 0.999998 | 0.999998 | 1 | 1.37841e-10 |
| **Hessen** | 26 | Chr1 | 5203996 | 5203997 | C | H26 10 | CC->CT | 0.999997 | 0.999997 | 1 | 1.37841e-10 |
| **Hessen** | 26 | Chr1 | 8612891 | 8612892 | T | H26 10 | TT->GT | 0.999919 | 0.999919 | 1 | 7.09277e-10 |
| **Hessen** | 26 | Chr1 | 21746287 | 21746288 | T | H26 4 | TT->AT | 1 | 1 | 1 | 7.0922e-10 |
| **Hessen** | 26 | Chr1 | 26975594 | 26975595 | T | H26 4 | TT->GG | 0.903972 | 0.903971 | 0.902162 | 1.41808e-11 |
| **Hessen** | 26 | Chr1 | 26975632 | 26975633 | A | H26 4 | AA->TT | 0.903972 | 0.903971 | 0.902162 | 1.41808e-11 |
| **Hessen** | 26 | Chr1 | 59736145 | 59736146 | C | H26 4 | CC->CT | 0.996931 | 0.996931 | 1 | 1.38265e-10 |
| **Hessen** | 26 | Chr2 | 31314215 | 31314216 | C | H26 4 | CC->CT | 1 | 1 | 1 | 1.37841e-10 |
| **Hessen** | 26 | Chr2 | 36518594 | 36518595 | A | H26 4 | AA->AG | 0.999999 | 0.999999 | 1 | 7.09221e-10 |
| **Hessen** | 26 | Chr1 | 13349686 | 13349687 | G | H26 6 | GG->GT | 0.999747 | 0.999747 | 1 | 1.37876e-10 |
| **Hessen** | 26 | Chr1 | 47932929 | 47932930 | G | H26 6 | GG->AG | 0.97538 | 0.97538 | 1 | 1.4132e-10 |
| **Hessen** | 26 | Chr1 | 52844991 | 52844992 | C | H26 6 | CC->TT | 1 | 1 | 0.99999 | 2.27582e-16 |
| **Hessen** | 26 | Chr2 | 15434190 | 15434191 | C | H26 6 | CC->CT | 0.999956 | 0.999956 | 1 | 1.37847e-10 |
| **Hessen** | 26 | Chr2 | 36579292 | 36579293 | C | H26 6 | CC->CT | 1 | 1 | 1 | 1.37841e-10 |
| **Hessen** | 26 | Chr3 | 23477115 | 23477116 | C | H26 6 | CC->CT | 1 | 1 | 1 | 1.37841e-10 |
| **Hessen** | 26 | Chr1 | 49977388 | 49977389 | T | H26 XX | TT->CT | 0.995752 | 0.995752 | 1 | 7.12246e-10 |
| **Hessen** | 26 | Chr2 | 4885896 | 4885897 | G | H26 XX | GG->GT | 0.999801 | 0.999801 | 1 | 1.37868e-10 |
| **Hessen** | 26 | Chr2 | 4885897 | 4885898 | A | H26 XX | AA->AT | 0.999842 | 0.999842 | 1 | 7.09332e-10 |
| **Hessen** | 26 | Chr2 | 55485182 | 55485183 | A | H26 XX | AA->AT | 0.962024 | 0.962024 | 1 | 7.37217e-10 |
| **Andalusia** | 15 | Chr1 | 9893906 | 9893907 | T | S15 12 | TT->AA | 0.988578 | 0.988578 | 0.945767 | 6.85645e-12 |
| **Andalusia** | 15 | Chr2 | 48529008 | 48529009 | T | S15 12 | TT->GG | 0.903972 | 0.903972 | 0.902162 | 1.41808e-11 |
| **Andalusia** | 15 | Chr2 | 12981217 | 12981218 | T | S15 21 | TT->AT | 0.987515 | 0.987515 | 1 | 7.18187e-10 |
| **Andalusia** | 15 | Chr2 | 11728539 | 11728540 | T | S15 21 | TT->AT | 0.934851 | 0.934851 | 1 | 7.58645e-10 |
| **Andalusia** | 15 | Chr2 | 12201837 | 12201838 | G | S15 21 | CC->CG | 1 | 1 | 1 | 1.0544e-14 |
| **Andalusia** | 15 | Chr2 | 28390820 | 28390821 | T | S15 22 | TT->CT | 0.917725 | 0.917725 | 0.999971 | 7.72803e-10 |
| **Andalusia** | 15 | Chr2 | 28390825 | 28390826 | C | S15 22 | CC->CT | 0.906033 | 0.906033 | 0.999259 | 1.52137e-10 |
| **Andalusia** | 15 | Chr1 | 19063562 | 19063563 | G | S15 25 | GG->AG | 1 | 1 | 1 | 1.37841e-10 |
| **Andalusia** | 15 | Chr1 | 21419796 | 21419797 | C | S15 25 | CC->AC | 0.999987 | 0.999987 | 1 | 1.37843e-10 |
| **Andalusia** | 15 | Chr2 | 41572584 | 41572585 | A | S15 25 | AA->AT | 0.998644 | 0.998644 | 1 | 7.10183e-10 |
| **Andalusia** | 15 | Chr1 | 22725223 | 22725224 | G | S15 25 | GG->AG | 0.999941 | 0.999941 | 1 | 1.37849e-10 |
| **Andalusia** | 15 | Chr1 | 23745544 | 23745545 | A | S15 26 | AA->AT | 0.99987 | 0.99987 | 1 | 7.09312e-10 |
| **Andalusia** | 15 | Chr1 | 54571763 | 54571764 | T | S15 27 | TT->CC | 0.903972 | 0.903972 | 0.902162 | 1.41808e-11 |
| **Andalusia** | 15 | Chr1 | 57503467 | 57503468 | T | S15 27 | TT->CC | 0.999969 | 0.999969 | 0.991172 | 1.05279e-12 |
| **Andalusia** | 15 | Chr3 | 35265163 | 35265164 | C | S15 27 | CC->CT | 0.97538 | 0.97538 | 1 | 1.4132e-10 |

| | | | | | | | | | | | |
|---|---|---|---|---|---|---|---|---|---|---|---|
| **Andalusia** | 15 | Chr4 | 4126488 | 4126489 | C | S15 27 | CC->CT | 0.984518 | 0.984518 | 1 | 1.40009e-10 |
| **Andalusia** | 15 | Chr1 | 39118300 | 39118301 | G | S15 29 | GG->AG | 0.999999 | 0.999999 | 1 | 1.37841e-10 |
| **Andalusia** | 15 | Chr2 | 47418116 | 47418117 | C | S15 29 | CC->TT | 0.996747 | 0.996747 | 0.970332 | 7.04721e-13 |
| **Andalusia** | 15 | Chr1 | 47726901 | 47726902 | A | S15 30 | AA->AC | 0.999954 | 0.999954 | 1 | 7.09253e-10 |
| **Andalusia** | 15 | Chr3 | 9232995 | 9232996 | C | S15 30 | CC->CT | 1 | 1 | 1 | 1.37841e-10 |
| **Andalusia** | 15 | Chr1 | 10176936 | 10176937 | T | S15 5 | TT->AT | 0.99998 | 0.99998 | 1 | 7.09234e-10 |
| **Andalusia** | 15 | Chr1 | 11830383 | 11830384 | A | S15 5 | AA->AT | 1 | 1 | 1 | 7.0922e-10 |
| **Andalusia** | 15 | Chr1 | 49975173 | 49975174 | T | S15 5 | TT->CT | 0.988223 | 0.988223 | 0.999204 | 7.17672e-10 |
| **Andalusia** | 15 | Chr3 | 8719967 | 8719968 | T | S15 5 | TT->AT | 1 | 1 | 1 | 7.0922e-10 |
| **Andalusia** | 15 | Chr1 | 4697315 | 4697316 | C | S15 7 | CC->AC | 0.999714 | 0.999714 | 1 | 1.37881e-10 |
| **Andalusia** | 15 | Chr3 | 52678531 | 52678532 | A | S15 7 | AA->AT | 0.99998 | 0.99998 | 1 | 7.09234e-10 |
| **Andalusia** | 17 | Chr1 | 31675350 | 31675351 | C | S17 1 | CC->TT | 0.948938 | 0.948938 | 0.920992 | 1.38826e-12 |
| **Andalusia** | 17 | Chr2 | 16145264 | 16145265 | A | S17 1 | AA->GG | 0.999799 | 0.999799 | 0.983829 | 1.94325e-12 |
| **Andalusia** | 17 | Chr2 | 16871178 | 16871179 | A | S17 1 | AA->GG | 0.999799 | 0.999799 | 0.983829 | 1.94325e-12 |
| **Andalusia** | 17 | Chr2 | 16871181 | 16871182 | G | S17 1 | GG->TT | 0.99993 | 0.99993 | 0.991172 | 2.04623e-13 |
| **Andalusia** | 17 | Chr2 | 16871183 | 16871184 | G | S17 1 | GG->CC | 0.99993 | 0.999928 | 0.991172 | 6.82761e-17 |
| **Andalusia** | 17 | Chr2 | 17926295 | 17926296 | C | S17 1 | CC->GG | 1 | 1 | 0.999173 | 1.90199e-14 |
| **Andalusia** | 17 | Chr3 | 22570048 | 22570049 | T | S17 1 | TT->CC | 1 | 1 | 0.930152 | 2.65932e-11 |
| **Andalusia** | 17 | Chr2 | 51553992 | 51553993 | C | S17 2 | CC->CT | 0.991298 | 0.991298 | 1 | 1.39051e-10 |
| **Andalusia** | 17 | Chr3 | 10949074 | 10949075 | G | S17 2 | GG->GT | 0.999989 | 0.999989 | 1 | 1.37843e-10 |
| **Andalusia** | 17 | Chr2 | 4863511 | 4863512 | A | S17 21 | AA->AT | 1 | 1 | 1 | 7.0922e-10 |
| **Andalusia** | 17 | Chr2 | 6551270 | 6551271 | T | S17 21 | TT->CT | 0.999963 | 0.999963 | 1 | 7.09246e-10 |
| **Andalusia** | 17 | Chr2 | 6551271 | 6551272 | T | S17 21 | TT->CT | 0.999812 | 0.999812 | 1 | 7.09353e-10 |
| **Andalusia** | 17 | Chr2 | 6551272 | 6551273 | T | S17 21 | TT->CT | 0.999963 | 0.999963 | 1 | 7.09246e-10 |
| **Andalusia** | 17 | Chr2 | 54110097 | 54110098 | G | S17 21 | GG->AG | 0.998636 | 0.998636 | 1 | 1.38029e-10 |
| **Andalusia** | 17 | Chr3 | 20224600 | 20224601 | A | S17 21 | AA->AG | 0.999998 | 0.999998 | 1 | 7.09222e-10 |
| **Andalusia** | 17 | Chr4 | 12351836 | 12351837 | G | S17 21 | GG->GT | 0.996469 | 0.996469 | 1 | 1.38329e-10 |
| **Andalusia** | 17 | Chr1 | 13592948 | 13592949 | A | S17 23 | AA->AT | 0.999966 | 0.999966 | 1 | 7.09245e-10 |
| **Andalusia** | 17 | Chr1 | 40873388 | 40873389 | T | S17 23 | TT->AA | 0.903972 | 0.903972 | 0.902162 | 1.41808e-11 |
| **Andalusia** | 17 | Chr2 | 19826132 | 19826133 | G | S17 25 | GG->AG | 1 | 1 | 1 | 1.37841e-10 |
| **Andalusia** | 17 | Chr2 | 38235021 | 38235022 | C | S17 25 | CC->CT | 0.999554 | 0.999554 | 1 | 1.37903e-10 |
| **Andalusia** | 17 | Chr2 | 38235334 | 38235335 | T | S17 25 | TT->CT | 0.999243 | 0.999243 | 1 | 7.09758e-10 |
| **Andalusia** | 17 | Chr3 | 10640403 | 10640404 | A | S17 25 | AA->GG | 0.903972 | 0.903972 | 0.902162 | 1.41808e-11 |
| **Andalusia** | 17 | Chr3 | 11800290 | 11800291 | G | S17 25 | GG->GT | 0.999976 | 0.999976 | 1 | 1.37844e-10 |
| **Andalusia** | 17 | Chr3 | 12129044 | 12129045 | C | S17 25 | CC->CG | 1 | 1 | 1 | 1.37841e-10 |
| **Andalusia** | 17 | Chr2 | 40508687 | 40508688 | A | S17 29 | AA->AC | 0.999757 | 0.999757 | 1 | 7.09393e-10 |
| **Andalusia** | 17 | Chr2 | 40508714 | 40508715 | A | S17 29 | AA->AC | 0.998206 | 0.998206 | 1 | 7.10495e-10 |
| **Andalusia** | 17 | Chr3 | 2995437 | 2995438 | C | S17 29 | CC->CT | 0.999358 | 0.999358 | 1 | 1.3793e-10 |
| **Andalusia** | 17 | Chr2 | 19090157 | 19090158 | A | S17 3 | AA->GG | 0.903972 | 0.903972 | 0.902162 | 1.41808e-11 |
| **Andalusia** | 17 | Chr2 | 39460892 | 39460893 | G | S17 3 | GG->AG | 0.99854 | 0.99854 | 1 | 1.38043e-10 |

| | | | | | | | | | | | |
|---|---|---|---|---|---|---|---|---|---|---|---|
| **Andalusia** | 17 | Chr3 | 2563984 | 2563985 | T | S17 3 | TT->CT | 0.931591 | 0.931591 | 1 | 7.613e-10 |
| **Andalusia** | 17 | Chr3 | 7057544 | 7057545 | G | S17 3 | GG->AG | 0.974027 | 0.974027 | 1 | 1.41517e-10 |
| **Andalusia** | 17 | Chr3 | 50492435 | 50492436 | C | S17 3 | CC->CG | 0.999048 | 0.999048 | 1 | 1.37972e-10 |
| **Andalusia** | 17 | Chr2 | 33689425 | 33689426 | C | S17 30 | CC->CT | 0.950216 | 0.950216 | 0.965032 | 1.45063e-10 |
| **Andalusia** | 17 | Chr3 | 16341037 | 16341038 | A | S17 30 | AA->TT | 0.903972 | 0.903972 | 0.902162 | 1.41808e-11 |
| **Andalusia** | 17 | Chr3 | 16341038 | 16341039 | C | S17 30 | CC->TT | 0.974464 | 0.974464 | 0.945767 | 1.35189e-12 |
| **Andalusia** | 17 | Chr4 | 3980035 | 3980036 | G | S17 30 | GG->AG | 0.999574 | 0.999574 | 1 | 1.379e-10 |
| **Andalusia** | 17 | Chr2 | 1291075 | 1291076 | G | S17 6 | GG->GT | 0.999999 | 0.999999 | 1 | 1.37841e-10 |
| **Andalusia** | 17 | Chr2 | 13265171 | 13265172 | G | S17 6 | GG->AG | 0.999999 | 0.999999 | 1 | 1.37841e-10 |
| **Andalusia** | 20 | Chr1 | 33899615 | 33899616 | A | S20 10 | AA->AG | 0.98865 | 0.98865 | 0.999866 | 7.17363e-10 |
| **Andalusia** | 20 | Chr1 | 33635875 | 33635876 | C | S20 10 | CC->CT | 0.906033 | 0.906033 | 0.999259 | 1.52137e-10 |
| **Andalusia** | 20 | Chr1 | 33635891 | 33635892 | A | S20 10 | AA->AT | 0.956271 | 0.956271 | 0.999259 | 7.41652e-10 |
| **Andalusia** | 20 | Chr1 | 52044653 | 52044654 | T | S20 10 | TT->AA | 0.903972 | 0.903972 | 0.902162 | 1.41808e-11 |
| **Andalusia** | 20 | Chr3 | 52658027 | 52658028 | A | S20 10 | AA->TT | 0.903972 | 0.903971 | 0.902162 | 1.41808e-11 |
| **Andalusia** | 20 | Chr4 | 11217283 | 11217284 | C | S20 10 | CC->TT | 1 | 1 | 1 | 2.46522e-18 |
| **Andalusia** | 20 | Chr3 | 3847131 | 3847132 | C | S20 8 | CC->TT | 1 | 1 | 1 | 1.53891e-18 |
| **Andalusia** | 20 | Chr3 | 45884759 | 45884760 | T | S20 8 | TT->AA | 0.988578 | 0.988578 | 0.945767 | 6.85645e-12 |
| **Andalusia** | 20 | Chr4 | 15637298 | 15637299 | T | S20 8 | TT->CT | 0.977421 | 0.977421 | 0.965032 | 7.25604e-10 |
| **Andalusia** | 23 | Chr1 | 1053307 | 1053308 | A | S23 22 | AA->AC | 0.999952 | 0.999952 | 1 | 7.09254e-10 |
| **Andalusia** | 23 | Chr1 | 16399294 | 16399295 | A | S23 22 | AA->AT | 0.999842 | 0.999842 | 1 | 7.09332e-10 |
| **Andalusia** | 23 | Chr1 | 59914857 | 59914858 | G | S23 22 | GG->AG | 0.999999 | 0.999999 | 1 | 1.37841e-10 |
| **Andalusia** | 23 | Chr1 | 22708034 | 22708035 | G | S23 23 | GG->AG | 1 | 1 | 1 | 1.37841e-10 |
| **Andalusia** | 23 | Chr2 | 2170697 | 2170698 | C | S23 23 | CC->CT | 0.995941 | 0.995941 | 1 | 1.38403e-10 |
| **Andalusia** | 23 | Chr2 | 3456535 | 3456536 | C | S23 23 | CC->CT | 0.988269 | 0.988269 | 1 | 1.39477e-10 |
| **Andalusia** | 23 | Chr2 | 6378702 | 6378703 | G | S23 23 | AA->AG | 0.999993 | 0.999993 | 1 | 5.42463e-14 |
| **Andalusia** | 23 | Chr2 | 8232279 | 8232280 | T | S23 23 | TT->AT | 0.977877 | 0.977877 | 1 | 7.25265e-10 |
| **Andalusia** | 23 | Chr2 | 55197031 | 55197032 | G | S23 23 | GG->AG | 0.999554 | 0.999554 | 1 | 1.37903e-10 |
| **Andalusia** | 23 | Chr3 | 31447685 | 31447686 | G | S23 23 | GG->AG | 0.999986 | 0.999986 | 1 | 1.37843e-10 |
| **Andalusia** | 23 | Chr3 | 37246241 | 37246242 | T | S23 23 | TT->AA | 0.903972 | 0.903972 | 0.902162 | 1.41808e-11 |
| **Andalusia** | 23 | Chr4 | 11198257 | 11198258 | A | S23 23 | AA->GG | 0.999799 | 0.999799 | 0.983829 | 1.94325e-12 |
| **Andalusia** | 23 | Chr3 | 11459225 | 11459226 | A | S23 25 | AA->AT | 0.994337 | 0.994337 | 0.978644 | 7.1326e-10 |
| **Andalusia** | 23 | Chr2 | 39197458 | 39197459 | G | S23 25 | GG->AG | 0.989408 | 0.989408 | 1 | 1.39317e-10 |
| **Andalusia** | 23 | Chr2 | 21327140 | 21327141 | T | S23 25 | AA->AT | 0.979385 | 0.979385 | 1 | 1.25618e-13 |
| **Andalusia** | 26 | Chr1 | 8726430 | 8726431 | C | S26 11 | CC->CT | 1 | 1 | 1 | 1.37841e-10 |
| **Andalusia** | 26 | Chr3 | 2589021 | 2589022 | T | S26 11 | TT->AT | 0.99648 | 0.99648 | 1 | 7.11725e-10 |
| **Andalusia** | 26 | Chr3 | 38664628 | 38664629 | T | S26 11 | TT->GG | 0.903972 | 0.903972 | 0.902162 | 1.41808e-11 |
| **Andalusia** | 26 | Chr3 | 40028443 | 40028444 | A | S26 11 | AA->GG | 0.903972 | 0.903972 | 0.902162 | 1.41808e-11 |
| **Andalusia** | 26 | Chr1 | 36955143 | 36955144 | G | S26 14 | GG->AG | 0.999706 | 0.999706 | 1 | 1.37882e-10 |
| **Andalusia** | 26 | Chr4 | 1124669 | 1124670 | A | S26 14 | AA->AT | 1 | 1 | 1 | 7.0922e-10 |
| **Andalusia** | 26 | Chr4 | 3456009 | 3456010 | T | S26 14 | TT->CT | 0.977421 | 0.977421 | 0.965032 | 7.25604e-10 |

| | | | | | | | | | | | |
|---|---|---|---|---|---|---|---|---|---|---|---|
| **Andalusia** | 26 | Chr4 | 5228924 | 5228925 | T | S26 14 | TT->CT | 1 | 1 | 0.919078 | 7.0922e-10 |
| **Andalusia** | 26 | Chr2 | 12235278 | 12235279 | A | S26 15 | AA->AG | 0.993114 | 0.993114 | 1 | 7.14138e-10 |
| **Andalusia** | 26 | Chr4 | 11630284 | 11630285 | G | S26 15 | GG->AG | 0.999999 | 0.999999 | 1 | 1.37841e-10 |
| **Andalusia** | 26 | Chr1 | 6238478 | 6238479 | G | S26 3 | GG->AG | 0.983302 | 0.983302 | 1 | 1.40182e-10 |
| **Andalusia** | 26 | Chr1 | 17285160 | 17285161 | G | S26 3 | GG->AG | 0.999358 | 0.999358 | 1 | 1.3793e-10 |
| **Andalusia** | 26 | Chr2 | 7624512 | 7624513 | T | S26 3 | TT->CT | 0.998972 | 0.998972 | 1 | 7.0995e-10 |
| **Andalusia** | 26 | Chr2 | 43963883 | 43963884 | T | S26 3 | TT->CT | 0.99998 | 0.99998 | 1 | 7.09234e-10 |
| **Andalusia** | 26 | Chr2 | 54062233 | 54062234 | A | S26 3 | AA->AG | 0.999757 | 0.999757 | 1 | 7.09393e-10 |
| **Andalusia** | 26 | Chr2 | 52932792 | 52932793 | A | S26 4 | AA->AG | 0.999997 | 0.999997 | 1 | 7.09222e-10 |
| **Andalusia** | 26 | Chr2 | 54044797 | 54044798 | G | S26 4 | GG->CG | 0.999976 | 0.999976 | 1 | 1.37844e-10 |
| **Andalusia** | 26 | Chr3 | 5808288 | 5808289 | T | S26 4 | TT->AA | 0.987547 | 0.987547 | 0.944781 | 6.8636e-12 |
| **Andalusia** | 26 | Chr3 | 6067621 | 6067622 | G | S26 4 | GG->GT | 0.986994 | 0.986994 | 1 | 1.39657e-10 |
| **Andalusia** | 26 | Chr1 | 57734407 | 57734408 | C | S26 6 | CC->GG | 1 | 1 | 0.999994 | 1.34541e-16 |
| **Andalusia** | 26 | Chr2 | 20065827 | 20065828 | G | S26 6 | GG->GT | 0.999999 | 0.999999 | 1 | 1.37841e-10 |
| **Andalusia** | 26 | Chr3 | 20136735 | 20136736 | C | S26 6 | CC->CG | 1 | 1 | 1 | 1.37841e-10 |
| **Andalusia** | 26 | Chr3 | 20681519 | 20681520 | G | S26 6 | GG->AG | 0.999987 | 0.999987 | 1 | 1.37843e-10 |
| **Andalusia** | 26 | Chr4 | 393171 | 393172 | A | S26 6 | AA->AG | 0.999999 | 0.999999 | 1 | 7.09221e-10 |
| **Andalusia** | 26 | Chr1 | 25710879 | 25710880 | C | S26 8 | CC->CT | 0.99999 | 0.99999 | 1 | 1.37842e-10 |
| **Andalusia** | 26 | Chr1 | 46376348 | 46376349 | G | S26 8 | GG->AG | 1 | 1 | 1 | 1.37841e-10 |
| **Andalusia** | 26 | Chr2 | 27570482 | 27570483 | A | S26 8 | AA->AC | 0.999897 | 0.999897 | 1 | 7.09293e-10 |
| **Andalusia** | 26 | Chr4 | 13378703 | 13378704 | C | S26 8 | CC->CT | 0.999358 | 0.999358 | 1 | 1.3793e-10 |
| **Andalusia** | 26 | Chr1 | 19043649 | 19043650 | A | S26 9 | AA->AT | 0.999996 | 0.999996 | 1 | 7.09223e-10 |
| **Andalusia** | 26 | Chr1 | 61024871 | 61024872 | C | S26 9 | CC->CT | 0.999996 | 0.999996 | 1 | 1.37842e-10 |
| **Andalusia** | 26 | Chr2 | 53049595 | 53049596 | A | S26 9 | AA->AG | 0.996539 | 0.996539 | 1 | 7.11683e-10 |
| **Andalusia** | 26 | Chr2 | 56344351 | 56344352 | G | S26 9 | GG->AA | 1 | 1 | 1 | 4.70811e-20 |

Supplemental Figure 1. Plot of modelled reaction norms with 95% confidence intervals from Bayesian inference.

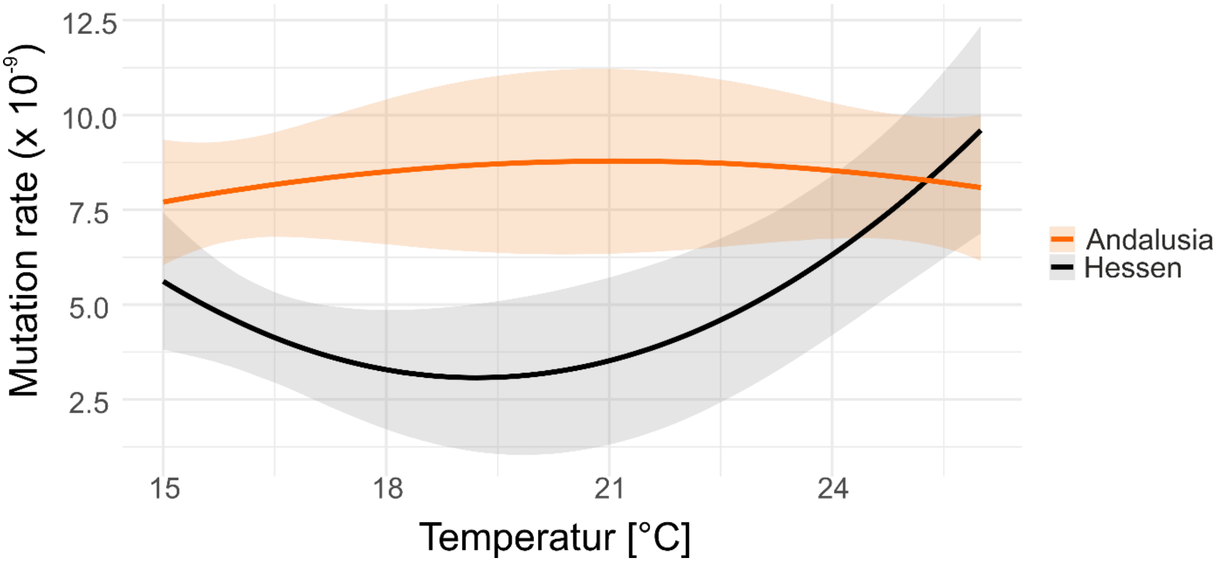
